# Bridging disciplines towards wastewater-based surveillance of antimicrobial resistance: frequency, local dynamics, and genomic characteristics of carbapenemase-producing *Klebsiella pneumoniae*

**DOI:** 10.64898/2026.03.02.707721

**Authors:** KA Werner, V Bajić, C Blumenscheit, D Baum, D Desirò, S Sedaghatjoo, J Barthelmes, AK Liebschner, C Förster, S Fuchs, SA Wolf, A Bethe, M Hölzer, B Walther

## Abstract

The World Health Organization (WHO) has designated carbapenemase-producing *Klebsiella pneumoniae* (CP-KP) as a critical priority pathogen due to its increasing importance for human health. As wastewater-based surveillance (WBS) is discussed as a complementary tool for classical systems regarding hazard forecasting and early-warning, we designed a “wet-lab to genomics” workflow to target CP-KP in raw influent wastewater samples to support method development processes across different scientific disciplines.

The CP-KP screening workflow was set up based on membrane filtration, selective chromogenic media for selective cultivation and the modified carbapenem inactivation method (mCIM) for confirming carbapenemase-production using 33 samples from four different wastewater treatment plants in North-Eastern Germany. All samples tested positive for CP-KP, with concentrations ranging between 10^2^ and 10^4^ colony-forming units (cfu) per 100 ml across the sample set. As a result, 320 isolates belonged to the *Klebsiella*, *Enterobacter*, *Citrobacter* (KEC)- group, with the majority being identified as KP (n= 297; 93%), including n= 253 (79%) verified CP-KP. Genotypic characterization of CP-KP by PCR revealed the predominance of *bla*_OXA-48_-related genes (n= 83) among isolates from all WWTPs.

As quality parameters, colony counts for viable *Escherichia coli* (EC) were employed as a proxy for valid wastewater samples and extended-spectrum beta-lactamase-producing *E. coli* (ESBL-EC) as indicator for AMR, with cfu/100 ml ranges from 10⁶ to 10⁷ and 10^2^ to 10⁴, respectively. To verify the screening outcome, a subset of 58 CP-KP from two WWTPs were subjected to whole genome sequencing (WGS). As a result, eight different sequence types (STs), i.e., ST147 and ST273 (both: clonal group 147), ST258, ST35, ST15, ST37, ST307, and ST485 were identified. These include clinically relevant STs clustering closest with fecal isolates from Germany when compared with Pathogenwatch-database entries. Moreover, WGS data enabled the identification of antibiotic resistance genes (ARGs), and the detection of closely related isolates within the WWTP dataset.

## 1. Introduction

According to the World Health Organization (WHO), “antimicrobial resistance (AMR) threatens the effective prevention and treatment of infectious diseases caused by bacteria, parasites, viruses and fungi” [1]. The WHO further declares that “AMR occurs when bacteria, viruses, fungi and parasites change over time and no longer respond to medicines making infections harder to treat and increasing the risk of disease spread, severe illness and death” [1]. Thus, AMR represents an acquired phenotypic trait of a pathogen, which is directly linked to the treatment of infectious diseases in humans, animals, and plants [2]. Recently, the revised European Urban Wastewater Treatment Directive (2024) integrated the One Health approach and defined AMR as “the ability of microorganisms to survive or to grow in the presence of a concentration of an antimicrobial agent which is usually sufficient to inhibit or kill microorganisms of the same species”. According to Article 17 of this directive, mandatory AMR monitoring in urban wastewater is required for agglomerations with 100,000 population equivalents or more in Europe [3].

Given the development of methods for wastewater-based surveillance (WBS) of AMR, it remains vital to acknowledge the natural evolution of antibiotics and antimicrobial resistance throughout the environment. Briefly, a wide range of antibiotics represent natural compounds, produced by environmental microorganisms, including - but not limited to - soil bacteria such as *Streptomyces* and other Actinomycetota, as well as multiple fungi (e.g., *Penicillium* spp.). Although their role within a specific habitat is still not fully understood [4], the general concept of producing antimicrobial compounds appears to represent a successful evolutionary trait for microorganisms, since these active metabolites have likely existed for millions of years, according to current estimates [5]. Antimicrobial resistance gene (ARG) evolution is closely tied to the natural biosynthesis of antibiotics, and “self-protection” motives of the producing cell have been discussed as a potential driver since the early 1970s [6]. However, many environmental bacteria possess intrinsic genetically determined resistance to different classes of antibiotics. The entirety of all intrinsic antibiotic resistances, i.e., the “intrinsic resistome”, is a naturally occurring phenomenon that predates antibiotic chemotherapy [7, 8]. The mobile resistome, on the other hand, encompasses ARGs that gained mobility via horizontal gene transfer [9], a process heavily influenced by anthropogenic activities [10-12]. ARGs conferring reduced susceptibility to antibiotics may be spread between bacteria inhabiting different ecosystems, encompassing environmental, commensal, and, last but not least, (facultative) pathogenic bacterial species that cause infectious diseases [13]. Consequently, monitoring AMR within urban wastewater, which represents a classical interface within the One Health continuum, depends on several key considerations. First, the target for AMR monitoring should be relevant to public health as defined by national and international health agencies [3, 14, 15]. Second, assessing AMR frequencies in terms of species exhibiting defined resistance phenotypes, e.g., carbapenemase-producing (CP) Enterobacterales, enables the identification of novel resistance mechanisms, variants, and combinations as well as sudden changes in resistance profiles [16-18]. Third, resident environmental microorganisms living in biofilm structures seem fully adapted to the challenges associated with the “wastewater system” habitat, including undulating nutrient accessibilities, the presence of selective agents (i.e., surfactants, biocides, antibiotics, heavy metals, pesticides, drugs and many others), variable oxygen and ammonia concentrations, as well as local temperature fluctuations. Consequently, resident (environmental) antibiotic resistant bacteria (ARB) such as *Pseudomonas* spp., *Acinetobacter* spp., and *Aeromonas* spp. harbor a broad range of mechanisms conferring tolerances and resistances, allowing for survival in these harsh conditions [19-21].

In recent years, WBS of AMR and especially its potential targets have been a subject of a broad scientific discussion, including intestinal bacteria such as Enterobacterales [16, 22-25], specifically *Escherichia coli* (EC) [26-29], and *Klebsiella pneumoniae* (KP) [26, 30, 31]. In this context, sequencing approaches such as whole-genome sequencing (WGS) of bacterial isolates and shotgun metagenomics are essential in supporting the detection and characterization of microbial species and ARGs from wastewater samples [32-37]. In particular, the isolation of bacterial species from wastewater samples, including their cultivation, sequencing, subsequent genomic reconstruction, and annotation can contribute to a better understanding of bacterial phenotype characteristics by providing molecular insights, including - but not limited to - the presence of resistance genes. On the other hand, metagenomic approaches may provide insights into the general population in wastewater and the broader resistome of the entire bacterial community within the wastewater system, in which sequencing depth, among other factors, is a particularly decisive parameter for detecting rare targets [38, 39].

Since bacteria of the human microbiota, as well as opportunistic pathogens capable of infecting humans, are typically well adapted to the conditions provided by warm-blooded hosts, their sheer survival in environments such as wastewater systems seems a considerable challenge [40-42]. In addition to chemical stressors, a wide array of microorganisms, many of which are specifically adapted to the wastewater environment, likely induce continuous changes in the bacterial composition of wastewater systems [40, 42-44]. Lower temperatures are likely to favor the proliferation of cold-adapted resident environmental bacteria, which frequently carry ARGs, such as *Pseudomonas* spp. (reviewed in [45]), *Aeromonas* spp. (reviewed in [46-48]), and *Acinetobacter* spp. (reviewed in [48-50]), or Gram-positive spore-forming bacteria (reviewed in [48]). Consequently, storage at 4 °C overnight prior to processing wastewater samples might introduce process-associated bias regarding detection rates of ARGs and human-associated bacteria (including antimicrobial resistant strains).

In 2021, the WHO Tricycle Protocol adopted the One Health approach using an AMR indicator organism, namely extended-spectrum beta-lactamase (ESBL)-producing *E. coli* (ESBL-EC), to streamline global surveillance efforts [27]. Likewise, monitoring of a single target seems feasible compared to an assortment of AMR pathogens and genes [51]. The importance of sensitive methods for targeting rare subpopulations such as carbapenemase-producing *K. pneumoniae* (CP-KP) was addressed only recently [30], and a systematic review on AMR wastewater surveillance declared clear and consistent reporting of study methods as a prerequisite to identify optimal practice [52].

The aim of this proof-of-principle study is to provide a “hands-on” procedure for the quantification of a specific AMR target in urban wastewater using a culture-based approach, combined with molecular methods, including subsequent WGS. This multidisciplinary approach provides insights into the diversity and resistance profiles of the respective targets. A step-by-step approach for the isolation and characterization of CP-KP is thus presented. It should be noted that a detailed epidemiological assessment of the WGS results is not the primary focus of this culture method-centered study. Here, we provide a robust and conclusive workflow for the detection of CP-KP, which is currently considered a suitable target for WBS due to its relevance to human health according to the WHO bacterial priority pathogens list [14, 15].

## 2. Material & Methods

### 2.1 Sampling

Between July and November 2024, raw influent wastewater samples were collected from four different wastewater treatment plants (WWTP) in North-Eastern Germany. The sampling process was recently described in detail [53]. Briefly, twenty-four-hour composite samples of primary clarified wastewater were transported *via* temperature-controlled shipment (4 °C) to the Microbiological Risks unit at the German Environment Agency (Berlin) and immediately further processed (within ten hours after sampling). Upon arrival, pH and temperature were measured, followed by sample homogenization *via* gentle inversion (overhead shaker reax20, Heidolph Scientific Products GmbH, Germany) for 15 minutes.

### 2.2 Pretesting: evaluation of two distinct temperature regimes for the detection of target colonies on selective chromogenic media

The study started with a pretest to identify the requirements for reliable KP detection in wastewater, a matrix that is, by nature, heavily loaded with non-target bacteria. Hence, a comparative analysis following previously published incubation regimes ([54] as cited in [16], and [55]) was conducted using influent wastewater samples collected throughout three consecutive weeks at two WWTPs. Briefly, wastewater-associated bacteria recovered by placement of loaded filters on selective media were subjected to either an incubation at 36 ± 2 °C for 18 ± 2h, or a 4-5h-incubation at 36 ± 1 °C followed by a 21 ± 3h incubation at 44 ± 0.5 °C. A serial dilution starting with 2 ml of influent wastewater was used as described in detail below and in Figure 1, with 0.5 and 1 ml wastewater for CP-KP detection as the primary target. Furthermore, 0.1 and 0.5 µl volumes were used to assess concentrations for EC, and 50 and 100 µl for ESBL-EC. Each volume was augmented with Phosphate-buffered saline (PBS) (Life Technologies Limited, Paisley, UK) to a final volume of 30 ml (pH ∼ 7.4) to achieve an appropriate volume for membrane filtration. Filtration was performed in triplicate for each target and wastewater volume investigated, yielding six selective plates per target and incubation regime for each sample. To test for significant differences between the incubation regimes, a two-tailed, paired Student’s t-test was conducted. Statistical significance was defined as a p-value below 0.05, p-values below 0.001 were considered highly significant. Results are provided in supplemental file 1.

**Figure 1.**
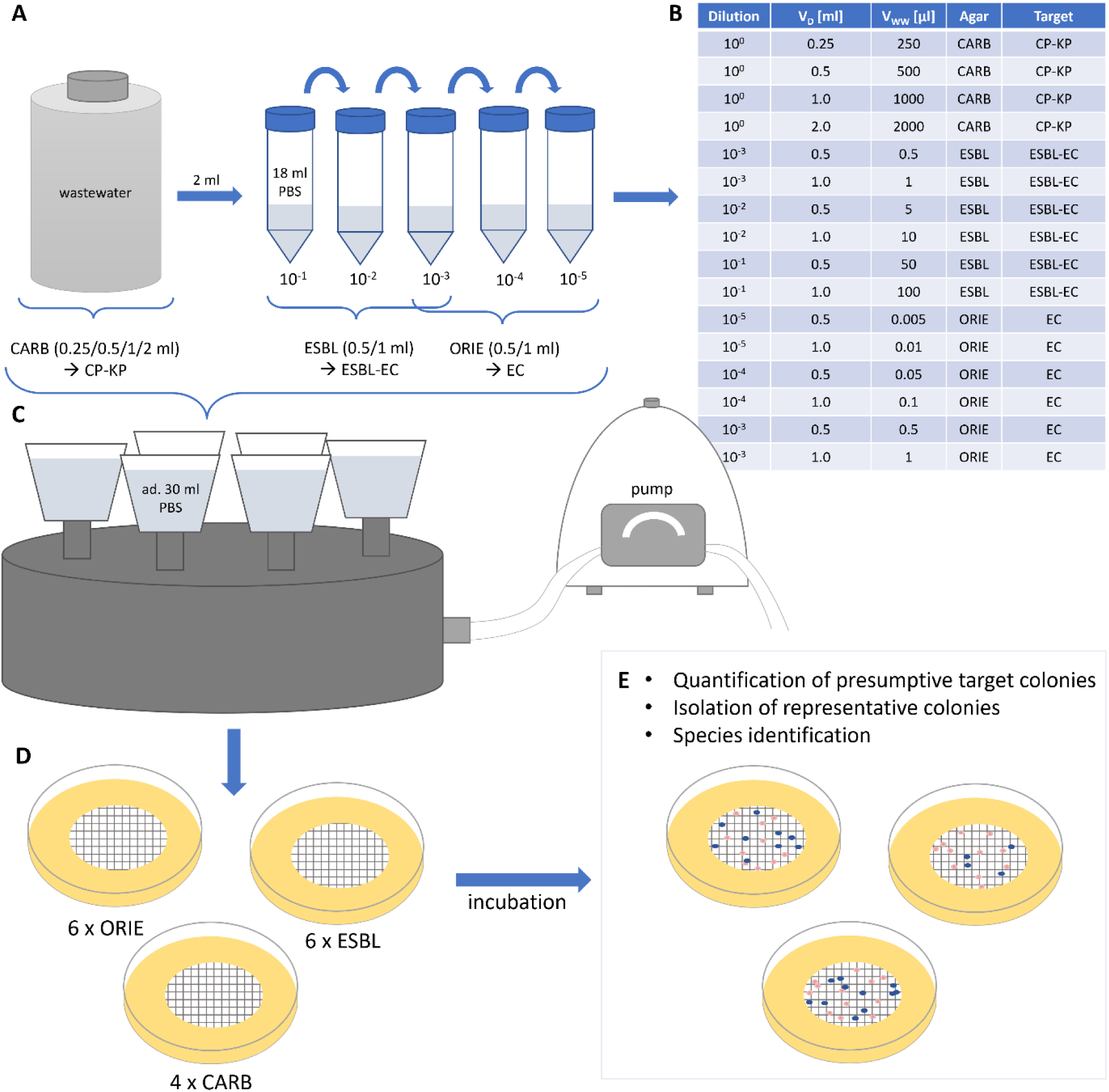
Graphical summary of the experimental procedure identifying CP-KP, EC, and ESBL-EC in influent wastewater. A serial dilution was prepared from influent wastewater using PBS (10^-1^ to 10^-^⁵) (A). Different volumes were filtered to detect presumptive CP-KEC, EC, and ESBL-EC (B). To ensure uniform bacterial distribution on the respective filters, volumes were supplemented with PBS to a final volume of 30 ml (C). Loaded filters were placed on chromogenic media as follows: CHROMagar^TM^ mSuperCARBA (CP-KP), CHROMagar^TM^ Orientation (EC), and CHROMagar^TM^ ESBL (ESBL-EC) (D). Following incubation, target colonies were quantified according to the manufacturer’s instructions. Presumptive CP-KEC grown on CHROMagar^TM^ mSuperCARBA were subjected to species identification and further characterization. Up to ten representative presumptive ESBL-EC colonies from at least two plates were used for species verification (E). Abbreviations: PBS, phosphate-buffered saline; CARB, CHROMagar^TM^ mSuperCARBA; ESBL, CHROMagar^TM^ ESBL; ORIE, CHROMagar^TM^ Orientation; CP-KP, carbapenemase-producing *K. pneumoniae*; ESBL-EC, extended spectrum beta lactamase *E. coli*; EC, *E. coli*; V_D_, filtrated volume of respective dilution; V_WW_, absolute volume of raw wastewater on filter.

### 2.3 Screening procedure for the target organism (CP-KP) and fecal indicators (EC, ESBL-EC)

A screening procedure was established, with KP positive for carbapenemase-production (CP) as the primary target (CP-KP; Figure 1). Samples representing four WWTPs were analyzed, with samples from WWTP 1 and 2 collected between week 31 and 48 of 2024 and samples from WWTP 3 and 4 collected between week 42 and 48 of 2024.

To identify CP-KP, 0.25, 0.5, 1, and 2 ml of wastewater were supplemented with PBS to a final volume of 30 ml. Subsequently, the samples were filtered using a SolarVac 601 MB filtration device (Rocker Scientific Co., Ltd., Taiwan) together with the Alligator 200 pump (Rocker Scientific Co., Ltd., Taiwan). The resulting four filters (cellulose nitrate filters 0.45 µm, Sartorius AG, Göttingen, Germany) were aseptically placed on mSuperCARBA (MAST Diagnostica GmbH, Reinfeld, Germany). Incubation at elevated temperatures was carried out as described above ([54] as cited in [16]).

According to the manufacturer’s guidelines, a metallic blue colony phenotype is a typical characteristic of Enterobacteria belonging to the group consisting of *Klebsiella* spp., *Enterobacter* spp. and *Citrobacter* spp. (“KEC”-group; Figure 1E). Consequently, colonies associated with a metallic blue appearance on mSuperCARBA plates were recorded as presumptive CP-KEC selected for further processing.

Presumptive CP-KEC were enumerated following the principles for colony selection stated in DIN EN ISO 8199:2008 [56]. In brief, 10 metallic blue colonies per sample from at least two different agar plates exhibiting the desired phenotype were subcultured on columbia blood agar plates (Life Technologies GmbH, Darmstadt, Germany). Up to seven target colonies were isolated from a single plate. Plates were excluded if bacterial overgrowth occurred or if more than 200 colonies resembling the typical target colony were present, as described before [16].

Two indicators were used to complement the CP-KEC assessment, EC and ESBL-EC following recommendations from the WHO in the “Tricycle protocol” [55]. EC- and ESBL-EC-screening was performed using CHROMagar^TM^ Orientation and CHROMagar^TM^ ESBL, respectively (MAST Diagnostica GmbH, Reinfeld, Germany). For each indicator, three dilution steps were utilized, containing different amounts of wastewater (Figure 1A). Per dilution step investigated, aliquots of 0.5 ml and 1 ml were supplemented with PBS to a final volume of 30 ml to facilitate membrane filtration using 0.45 µm cellulose nitrate filters. Thus, the final wastewater volumes analyzed were 0.005, 0.01, 0.05, 0.1, 0.5, and 1 µl for EC, and 0.5, 1, 5, 10, 50, and 100 µl for ESBL-EC. Additional details are provided in Figure 1B.

To verify species identity, presumptive CP-KEC (including CP-KP) and ESBL-EC were investigated by use of MALDI-TOF MS (Biotyper® Sirius, Bruker Daltronics GmbH & Co. KG, Bruker Corporation, Massachusetts, USA) in combination with flexControl (version 3.4) and MBT Compass HT software programs (version 5.1.300) along with the MBT Compass Library (version 12.0.0.0) and the BTyp 2.0-Sec-Library (version 1.0) according to the manufacturer’s instructions (Bruker Daltronics GmbH & Co. KG, Bruker Corporation, Massachusetts, USA).

To confirm the species identity of presumptive EC from CHROMagar^TM^ Orientation, presumptive EC colonies collected during an eight-week sampling period were initially identified using MALDI-TOF MS. To save costs, the error rate of 0.8% (2/240) on CHROMagar^TM^ Orientation was considered negligible for EC ratio calculations as fecal indicator and subsequent identification of EC was limited to colony phenotype appearances.

### 2.4 Antimicrobial susceptibility testing of KP

Verified KP isolates were subjected to antibiotic susceptibility testing (AST) using a VITEK 2 COMPACT system (Biomérieux, Marcy l’Étoile, France), providing minimum inhibitory concentrations (MICs) for selected antibiotics representing cephalosporines (e.g., cefotaxime), carbapenems (e.g., imipenem, meropenem), chinolones (ciprofloxacin), aminoglycosides (gentamicin), and sulfonamides (trimethoprim-sulfamethoxazole). Further characterization included phenotypic confirmation of i) carbapenemase-production of KP by use of the modified carbapenem inactivation method (mCIM) and ii) ESBL-production confirmation according to CLSI guidelines (CLSI, https://clsi.org/).

### 2.5 Molecular determination of carbapenem resistance genes by use of PCR screening

PCR initially was utilized to screen all CP-KP for the presence of common carbapenemase-encoding genes (i.e., *bla*_OXA-48_-related genes*, bla*_NDM_*, bla*_KPC_*, bla*_VIM_ and *bla*_IMP_) as previously described (details are provided in Table S1) [57-59].

### 2.6 Correction of wet-lab screening data

In accordance with the procedures outlined in EN ISO 8199:2018, enumeration of the target CP-KP was performed using the number of presumptive CP-KEC colonies grown on mSuperCARBA, followed by a correction based on the results obtained by MALDI-TOF species verification and mCIM-confirmation of carbapenemase production for the representative selection of subcultures.

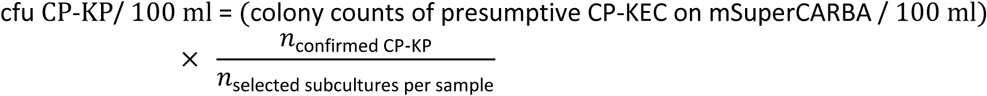

For ESBL-EC, cfu values were adjusted based on the proportion of subcultured colonies confirmed as EC.

### 2.7 DNA extraction and quantification for whole-genome sequencing

Genomic DNA (gDNA) of 58 selected CP-KP isolates was extracted using the Monarch® Genomic DNA Purification Kit (New England Biolabs GmbH, Frankfurt am Main, Germany) according to the manufacturer’s instructions for gram-negative bacteria. DNA was quantified using a Qubit 4 Fluorometer and the Qubit^TM^ 1x dsDNA HS Assay Kit (both invitrogen by ThermoFisher Scientific, Massachusetts, USA), and stored at −20 °C until use.

### 2.8 Library preparation and whole genome sequencing

Dual-indexed or unique dual-indexed libraries were prepared from 1 ng of genomic DNA using the Nextera XT DNA Library Preparation Kit (Illumina, San Diego, CA, USA), following the manufacturer’s protocol with all reagent volumes scaled down to half. The library preparation procedure was fully automated on the Hamilton Microlab STAR liquid handling system (Hamilton company, Germany). Briefly, the gDNA was initially fragmented and tagmented in a single enzymatic step, followed by PCR amplification to enrich adapter-ligated fragments and incorporate index sequences. Library cleanup was performed using the MagSi-NGSPREP Plus magnetic beads (Steinbrenner Laborsysteme GmbH, Germany). The prepared libraries were quantified using the QuantiFluor® dsDNA System Kit (Promega) on an Agilent BioTek Synergy HTX multi-mode microplate reader. Fragment size distribution was assessed using the 5300 Fragment Analyzer system (Agilent Technologies, Malaysia) with the High Sensitivity NGS Fragment Kit (Agilent, CA, USA).

The final libraries were normalized to 2 nM, pooled equimolarly, diluted to the recommended loading concentration, and sequenced on the Illumina NextSeq 2000 platform (Illumina Inc., Singapore) using v3 reagents with a 2×150 bp read configuration, following the manufacturer’s instructions. The obtained coverage was >150X with an average read length of 144 bp. Illumina raw read data sequenced for this study is available on the Sequence Read Archive (SRA) of the National Center for Biotechnology Information (NCBI) under Bioproject ID PRJEB107251.

### 2.9 Genomic characterization and comparison with publicly available data

Genome assembly and quality control were performed using the Generic Assembly and Reconstruction (GARI) pipeline (v1.0.0; https://github.com/rki-mf1/GARI). GARI employs *fastp* (v0.23.4; [60]) for adapter sequence removal and assessment of read quality statistics, and Shovill (v1.1.0; https://github.com/tseemann/shovill) for *de novo* genome assembly. The generated assemblies were evaluated based on multiple assembly statistics such as average coverage, assembly length, completeness, and contamination (see GARI pipeline results for a detailed overview) (Table S3).

Assembled genomes in FASTA format were uploaded to the Pathogenwatch web application (v23.1.5), which integrates several community-driven tools for genomic antimicrobial resistance (AMR) prediction, multilocus sequence typing (MLST), core genome MLST (cgMLST) typing, and phylogenetic assessment.

To contextualize the wastewater isolates, we compared them with publicly available *Klebsiella pneumoniae* genomes on Pathogenwatch (v23.1.5) as of February 20, 2025. After determining that the wastewater isolates encompassed eight different sequence types (STs), we created multiple collections on the Pathogenwatch web-based platform containing all publicly available genomes corresponding to one of those eight STs (Table S2). From these Pathogenwatch collections, we downloaded the hashed cgMLST profiles and merged them into a single dataset comprising 14,590 publicly available *Klebsiella pneumoniae* genomes belonging to the same eight STs as the wastewater samples.

The combined cgMLST profile dataset was then utilized as input for a modified version of the cgmlst-dists tool (v0.4.0; https://github.com/KHajji/cgmlst-dists/tree/master), which calculates pairwise distances using hashed profiles (via the “-H” parameter), enabling further analysis without strictly requiring numeric allele names.

For 58 wastewater isolates, a neighbor-joining tree was constructed in Pathogenwatch based on pairwise single-nucleotide polymorphism (SNP) distances across a curated set of 1,972 core genome loci [61]. The resulting tree was exported in Newick format for visualization in R (v4.3.0; [62]), alongside the genomic AMR prediction output from Pathogenwatch. For data wrangling, the *tidyverse* package (v2.0.0; [63]) was utilized. For visualization of the phylogenetic tree and heatmaps we used *treeio* (v1.26.0; [64]) and *tidyheatmaps* (v0.2.1; [65]), while additional plots were generated using *ggplot2* (v3.5.1; [66]).

### 2.10 Snippy analyses

To increase the resolution for the phylogenetic assessment of the wastewater isolates, we performed SNP-based analysis on assemblies using snippy (v4.6.0; https://github.com/tseemann/snippy). For each ST represented by at least two isolates, we selected the sample with the best assembly quality (based on metrics including N50, percentage of complete single-copy BUSCOs, percentage of reads mapped back to assembly, and contamination; see Table S3) as the genomic reference sequence. All other isolate assemblies belonging to the same ST were then aligned and compared against this reference using snippy, enabling identification of core SNPs within each ST. Pairwise SNP distances were then visualized in R using the *tidyheatmaps* package.

## 3. Results

### 3.1 Description of the sample set

In total, 33 influent wastewater samples (i.e., 11 samples from WWTP 1, 10 from WWTP 2, 6 from WWTP 3, and 6 from WWTP 4) were used to set up the general procedure. The performance of the incubation regimen was evaluated during the pretesting phase; the results are provided in supplementary chapter 1. The subsequent workflow, as described in Fig. 1, resulted in 325 presumptive CP-KEC. Amongst these, 320 isolates belonged to the KEC group, with the majority being identified as KP (297 of 320; 93%), including 253 (79%) verified CP-KP.

### 3.2 AMR concentration: screening results for target CP-KP, and indicators EC and ESBL-EC CP-KP concentrations in wastewater samples

As a result, concentrations of presumptive CP-KEC were, with few exceptions, in overall agreement with the corresponding values obtained for verified CP-KP across all samples and WWTPs (Figure 2), particularly with respect to their overall order of magnitude, with colony counts ranging from 10² to 10⁴ cfu/100 ml (Figure 2).

**Figure 2.**
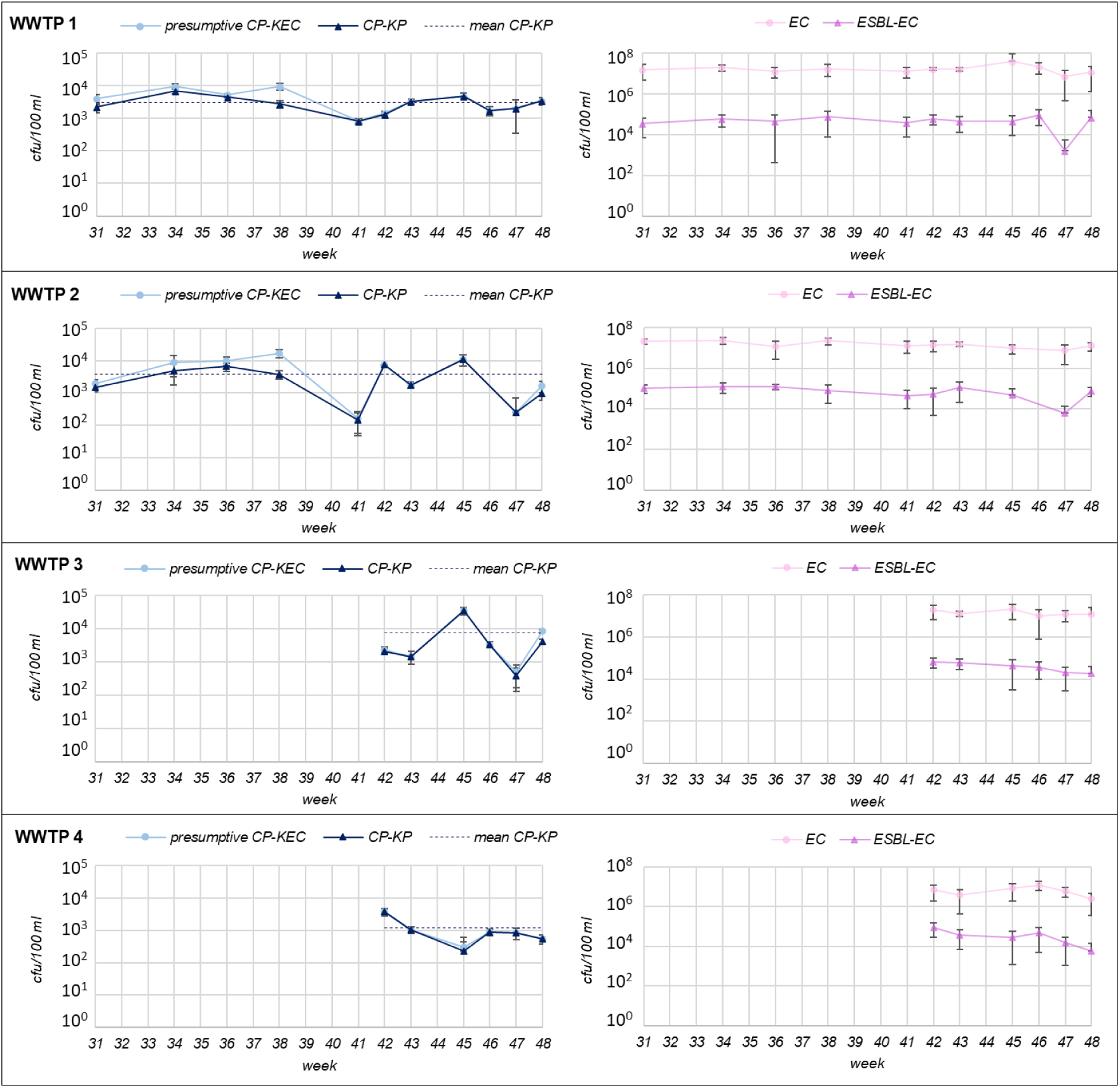
Temporal variation of presumptive CP-KEC, CP-KP, EC and ESBL-EC (cfu/100 ml) in wastewater samples. Temporal variation of target CP-KP, presumptive CP-KEC colony counts (left), and indicators (fecal & AMR) EC and ESBL-EC (right) across four WWTPs in 2024. Colony counts are depicted per 100 ml influent wastewater and represent mean values ± standard deviation (error bars) for each sampling point. Dotted lines indicate the mean CP-KP cfu/100 ml across the respective sample set (weeks 31-48/2024 for WWTP 1 and 2; weeks 42-48/2024 for WWTP 3 and 4). Abbreviations: WWTP, wastewater treatment plant; CP, carbapenemase-producing; KEC, *Klebsiella*/ *Enterobacter*/ *Citrobacter* group; KP, *Klebsiella pneumoniae*; EC, *Escherichia coli*; ESBL-EC, extended-spectrum beta-lactamase-producing *E. coli*; cfu, colony-forming units.

Mean cfu/100 ml values obtained for CP-KP were 3.07 x 10^3^ cfu/100 ml (WWTP 1), 3.79 x 10^3^ cfu/100 ml (WWTP 2), 7.66 x 10^3^ cfu/100 ml (WWTP 3), and 1.22 x 10^3^ cfu/100 ml (WWTP 4). Spatio-temporal variation of CP-KP cfu counts was observed for the WWTPs investigated in this study. Exemplarily, CP-KP colony counts for week 45 exhibited a two-log_10_ difference, ranging from 5 x 10^2^ cfu/100 ml (minimum: WWTP 4) to 5 x 10^4^ cfu/100 ml (maximum: WWTP 3) (Figure 2). The minimum CP-KP counts were obtained for samples collected in week 41 (1.44 × 10^2^ cfu/100 ml, WWTP 2) and the maximum was recorded in week 45 (3.44 × 10^4^ cfu/100 ml, WWTP 3). In WWTP 3, CP-KP counts were comparatively high in week 45, with cfu’s within a similar order of magnitude as ESBL-EC (10^4^ cfu/100 ml).

#### EC and ESBL-EC concentrations in wastewater samples

Colony counts of EC as fecal indicators were temporally stable across all sampling weeks regarding their orders of magnitude. Colony counts ranged between 10^6^ and 10^7^ cfu/100 ml with mean values of 1.7 x 10^7^ cfu/100 ml (WWTP 1), 1.5 x 10^7^ cfu/100 ml (WWTP 2), 1.5 x 10^7^ cfu/100 ml (WWTP 3), and 6.6 x 10^6^ cfu/100 ml (WWTP 4). The minimum EC colony counts were obtained for samples collected in week 48 (2.47 × 10^6^ cfu/100 ml, WWTP 4), peaking counts were recorded in week 45 (3.73 × 10^7^ cfu/100 ml, WWTP 1) (Figure 2).

As an indicator of viable AMR bacteria, ESBL-EC illustrated roughly 100-fold lower colony counts compared to those of EC (Figure 3), with few deviations. For ESBL-EC, mean cfu/100 ml counts were 5.2 x 10^4^, 7.8 x 10^4^, 4.2 x 10^4^, 3.6 x 10^4^ for WWTP 1-4, respectively. The lowest colony counts were obtained for samples collected in week 47 (1.67 × 10^3^ cfu/100 ml, WWTP 1), and the highest counts were recorded in week 34 (1.28 × 10^5^ cfu/100 ml, WWTP 2) (Figure 2).

**Figure 3.**
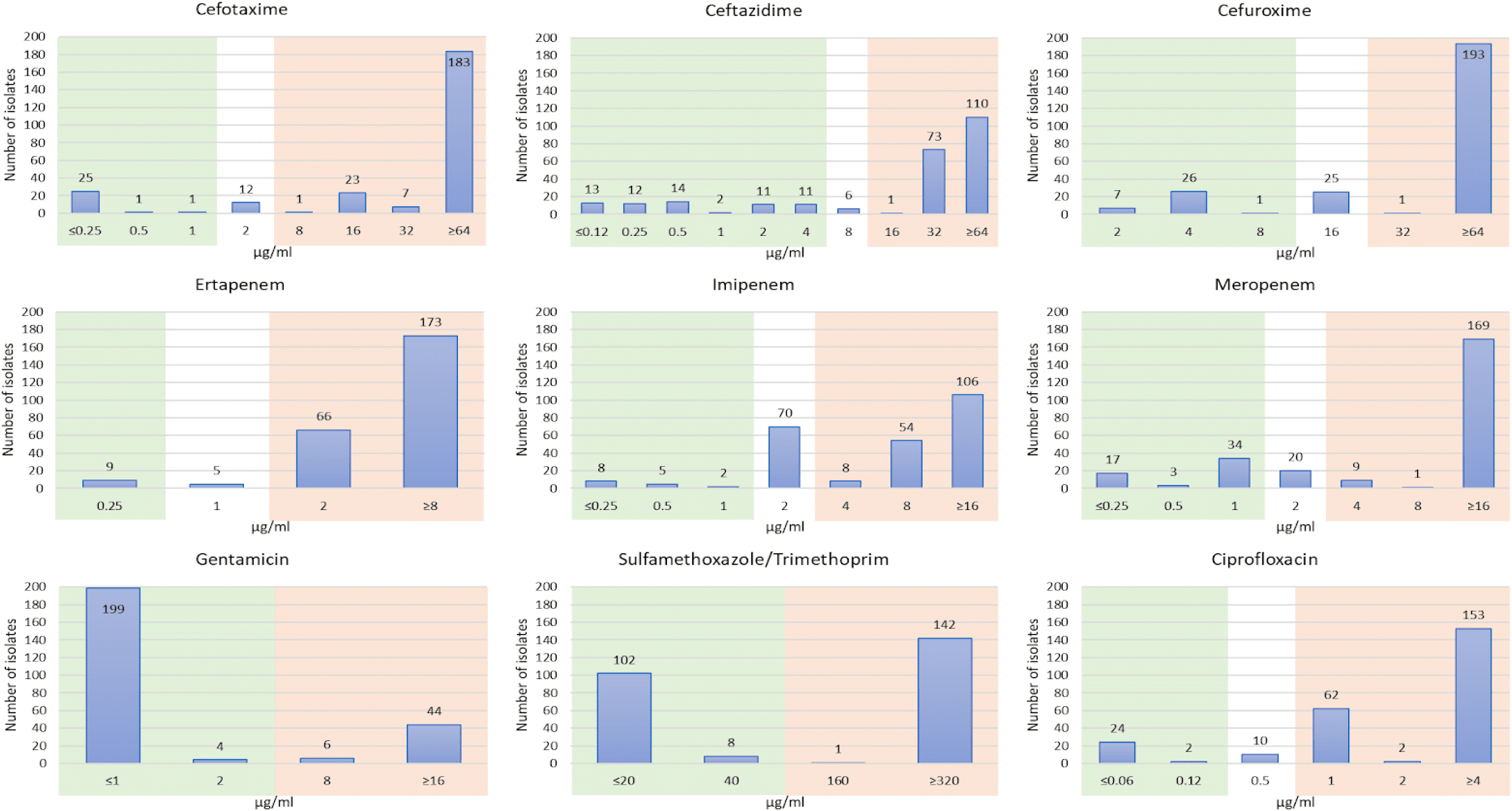
Distribution of MIC values for 253 CP-KP isolates across selected antimicrobial substances. The green-shaded areas of the graph indicate that, according to CLSI guidelines M100Ed35:2025, isolates would be classified as susceptible if isolated from human clinical cases. In contrast, the red-shaded areas denote the range at which clinical isolates would be classified as resistant for at least one specific indication (e.g., wound infection, pneumonia, blood stream infection).

### 3.3 CP-KP: antibiotic susceptibility profiles and PCR results

Overall, each of the wastewater samples investigated was positive for CP-KP (Table 1). AST results (MICs) were assessed for all subcultured CP-KP isolates as presented in Figure 3.

**Table 1.** CP-KP screening results obtained for 33 wastewater samples from four WWTPs.

| WWTP | week (2024) | sub-cultures (pheno-type) | confirmed KEC | confirmed KP | CP-KP | WGS | ESBL positive | PCR screening results |  |  |  |  |  |  |  |
| --- | --- | --- | --- | --- | --- | --- | --- | --- | --- | --- | --- | --- | --- | --- | --- |
|  |  |  |  |  |  |  |  | *bla*<sub>OXA-48</sub>-related genes | *bla*<sub>VIM</sub> | *bla*<sub>NDM</sub> | *bla*<sub>KPC</sub> | *bla*<sub>OXA-48</sub>-related genes + *bla*<sub>VIM</sub> | *bla*<sub>OXA-48</sub>-related genes+ *bla*<sub>NDM</sub> | *bla*<sub>OXA-48</sub>-related genes + *bla*<sub>KPC</sub> | *bla*<sub>NDM</sub> + *bla*<sub>KPC</sub> |
| 1 | 31 | 10 | 10 | 9 | 5 | 3 | 0 | 4 | 0 | 0 | 0 | 0 | 0 | 0 | 0 |
|  | 34 | 10 | 10 | 8 | 6 | 6 | 0 | 2 | 0 | 0 | 3 | 0 | 0 | 0 | 0 |
|  | 36 | 10 | 10 | 9 | 8 | 7 | 3 | 5 | 0 | 0 | 3 | 0 | 0 | 0 | 0 |
|  | 38 | 10 | 10 | 7 | 2 | 2 | 0 | 0 | 0 | 0 | 2 | 0 | 0 | 0 | 0 |
|  | 41 | 10 | 10 | 10 | 10 | 4 | 4 | 3 | 0 | 0 | 7 | 0 | 0 | 0 | 0 |
|  | 42 | 10 | 10 | 10 | 9 | 4 | 1 | 1 | 0 | 0 | 6 | 0 | 0 | 0 | 0 |
|  | 43 | 10 | 10 | 10 | 10 | 5 | 0 | 0 | 0 | 0 | 3 | 0 | 0 | 0 | 2 |
|  | 45 | 10 | 8 | 8 | 8 | 0 | 3 | 7 | 0 | 0 | 1 | 0 | 0 | 0 | 0 |
|  | 46 | 10 | 10 | 6 | 6 | 0 | 0 | 4 | 0 | 0 | 2 | 0 | 0 | 0 | 0 |
|  | 47 | 10 | 10 | 10 | 10 | 0 | 0 | 1 | 0 | 0 | 8 | 0 | 0 | 0 | 0 |
|  | 48 | 10 | 10 | 10 | 10 | 0 | 4 | 1 | 0 | 0 | 9 | 0 | 0 | 0 | 0 |
| 2 | 31 | 10 | 10 | 9 | 7 | 7 | 3 | 3 | 0 | 0 | 0 | 0 | 4 | 0 | 0 |
|  | 34 | 10 | 10 | 9 | 5 | 5 | 1 | 1 | 0 | 4 | 0 | 0 | 0 | 0 | 0 |
|  | 36 | 10 | 9 | 9 | 6 | 6 | 6 | 3 | 0 | 3 | 0 | 0 | 0 | 0 | 0 |
|  | 38 | 10 | 10 | 9 | 2 | 2 | 0 | 0 | 0 | 2 | 0 | 0 | 0 | 0 | 0 |
|  | 41 | 9* | 9 | 9 | 8 | 3 | 7 | 2 | 0 | 6 | 0 | 0 | 1 | 0 | 0 |
|  | 42 | 10 | 10 | 10 | 10 | 0 | 0 | 0 | 0 | 9 | 0 | 0 | 0 | 0 | 0 |
|  | 43 | 10 | 10 | 10 | 10 | 4 | 2 | 2 | 0 | 8 | 0 | 0 | 0 | 0 | 0 |
|  | 45 | 9 | 9 | 9 | 9 | 0 | 0 | 0 | 0 | 9 | 0 | 0 | 0 | 0 | 0 |
|  | 47 | 5* | 5 | 5 | 5 | 0 | 0 | 1 | 0 | 4 | 0 | 0 | 0 | 0 | 0 |
|  | 48 | 10 | 10 | 10 | 6 | 0 | 3 | 0 | 0 | 5 | 0 | 0 | 1 | 0 | 0 |
| 3 | 42 | 11 | 11 | 9 | 8 |  | 3 | 1 | 0 | 1 | 2 | 1 | 1 | 1 | 0 |
|  | 43 | 10 | 10 | 8 | 8 |  | 2 | 2 | 0 | 1 | 0 | 0 | 2 | 0 | 0 |
|  | 45 | 10 | 10 | 10 | 10 |  | 9 | 10 | 0 | 0 | 0 | 0 | 0 | 0 | 0 |
|  | 46 | 10 | 10 | 10 | 10 |  | 8 | 9 | 1 | 0 | 0 | 0 | 0 | 0 | 0 |
|  | 47 | 10 | 9 | 9 | 7 |  | 1 | 3 | 0 | 3 | 1 | 0 | 0 | 0 | 0 |
|  | 48 | 10 | 10 | 10 | 5 |  | 3 | 5 | 0 | 0 | 0 | 0 | 0 | 0 | 0 |
| 4 | 42 | 10 | 10 | 10 | 10 |  | 6 | 2 | 0 | 4 | 1 | 0 | 1 | 0 | 0 |
|  | 43 | 10 | 10 | 10 | 10 |  | 1 | 2 | 0 | 0 | 0 | 0 | 8 | 0 | 0 |
|  | 45 | 11 | 11 | 8 | 6 |  | 2 | 2 | 0 | 0 | 0 | 0 | 2 | 0 | 0 |
|  | 46 | 10 | 10 | 10 | 10 |  | 1 | 1 | 0 | 0 | 1 | 0 | 8 | 0 | 0 |
|  | 47 | 10 | 10 | 10 | 10 |  | 0 | 2 | 0 | 1 | 0 | 0 | 7 | 0 | 0 |
|  | 48 | 10 | 9 | 7 | 7 |  | 0 | 5 | 0 | 0 | 0 | 0 | 2 | 0 | 0 |

Information on samples, sampling period, subcultures, presumptive and confirmed CP-KP and PCR-screening results for four common carbapenemase-encoding genes. Grey areas: no data available. Asterisks (\*) indicate samples with a maximum number of presumptive CP-KEC colonies below ten. Abbreviations: WWTP, wastewater treatment plant; WGS, whole genome sequencing; KEC, *Klebsiella-Enterobacter-Citrobacter* group; KP, *Klebsiella pneumoniae*; CP-KP, carbapenemase-producing *Klebsiella pneumoniae* (species- and carbapenemase production verified); ESBL, extended-spectrum beta-lactamase.

Phenotypic AST was performed using antimicrobial substances from different antimicrobial families, including cephalosporines (cefotaxime, ceftazidime, cefuroxime), carbapenems (ertapenem, imipenem, meropenem), aminoglycosides (gentamicin), quinolones (ciprofloxacin), and folate pathway inhibitors (sulfamethoxazole/trimethoprim). Details on the distribution of MICs obtained for 253 CP-KP detected are given in Figure 3. AST profiles for each isolate are summarized in Table S4.

Genotypic characterization of CP-KP by PCR revealed the predominance of *bla*_OXA-48_-related genes (n= 83) among isolates from all WWTPs, followed by *bla*_NDM_ (n= 60) in WWTPs 2, 3 and 4, *bla*_KPC_ (n= 49) in WWTP 1, 3 and 4, and *bla*_VIM_ (n= 1) in WWTP 3, while *bla*_IMP_ was not detected. Co-occurrence of carbapenemase-encoding genes was observed in 41 isolates with *bla*_OXA-48_-related genes /*bla*_NDM-1_ (n= 37; WWTPs 2, 3 and 4) being the most common combination, followed by *bla*_NDM_ /*bla*_KPC_ (n= 2; WWTP 1), *bla*_OXA-48_-related genes /*bla*_VIM_ (n= 1; WWTP 3) and *bla*_OXA-48_-related genes /*bla*_KPC_ (n= 1; WWTP 3). Individual data for each isolate is provided in Table S4. Due to the limited panel of PCR-targeted genes, 19 isolates lacked a positive PCR result for a specific carbapenemase-encoding gene (Table 1). Further characterization using WGS revealed carbapenemase-encoding genes (i.e., *bla*_KPC-2_, *bla*_NDM-1_, *bla*_NDM-5_, *bla*_OXA-48_, *bla*_OXA-181_, *bla*_OXA-427_) in each of the CP-KP isolates sequenced (Table S4).

Fourteen CP-KP isolates recovered from samples of three WWTPs exhibited ertapenem, imipenem, and meropenem MICs below the CLSI-recommended thresholds for further testing for carbapenemase production (Figure 3 and Table S4). While PCR analysis revealed the presence of *bla*_OXA-48_-related genes in 13 of these, the evaluation of whole-genome data further revealed that the isolate lacking a specific PCR signal harbored the *bla*_OXA-48_ gene *bla*_OXA-181_ (Table S4).

### 3.4 Genomic profiling of CP-KP wastewater isolates and comparison to public genomic data

To gain further insights into the genetic makeup of CP-KP isolated from wastewater samples, 58 isolates from WWTP 1 and 2 were selected for WGS, following a stepwise approach. The initial sequencing batch covering isolates from samples representing week 31, 34, 36, and 38 accounted for the majority of CP-KP isolates (n=41). These isolates were selected to verify the sensitivity of the wet-lab screening process regarding CP-KP harboring carbapenemase-encoding genes (positive rate: 100%), as the PCR results (see Table S4) were inconclusive due to limitations of the screening panel. Isolates for the second sequencing batch following week 41 were selected to reveal the phylogenetic background of CP-KP exhibiting divergent AST results obtained from a specific wastewater sample, while most presumptive isogenic isolates (i.e., isolates sharing a similar phenotype and AST profile) were mostly excluded to save resources. An overview of isolate IDs, STs, and ARGs detected from WGS per week across the 12-week sampling period is provided in Table S4.

CP-KP belonging to different phylogenetic backgrounds were detected in most samples, including four different STs in WWTP 1 (weeks 36 and 42). Closely related genomes of isolates belonging to ST35 carrying *bla*_OXA-181_ were detected in samples of both WWTPs (Figure 4). Further isolates obtained from samples of WWTP 1 were assigned as follows: ST37 (n= 10, *bla*_KPC-2_), ST258 (n= 5, *bla*_KPC-2_), ST273 (n= 1, *bla*_KCP-2_), and ST485 (n= 10, *bla*_OXA-181_). Isolates assigned to ST15 (n= 2, both isolates: *bla*_OXA-48_ and *bla*_OXA-427_, one isolate in combination with *bla*_NDM-1_), and ST307 (n= 4, *bla*_NDM-1_) were obtained from WWTP 2 samples. In addition, WGS analysis of isolates belonging to ST147 (n= 14) highlighted two genomic clusters. In the first cluster, four out of six isolates were found to carry *bla*_OXA-48_ and *bla*_NDM-5_, while 8 isolates from the second cluster were associated with *bla*_NDM-1_ (Figure 4). WGS analyses revealed between one and three (isolate ID 8205) carbapenemase-encoding genes for KP-CP (Figure 4 and Table S4).

**Figure 4.**
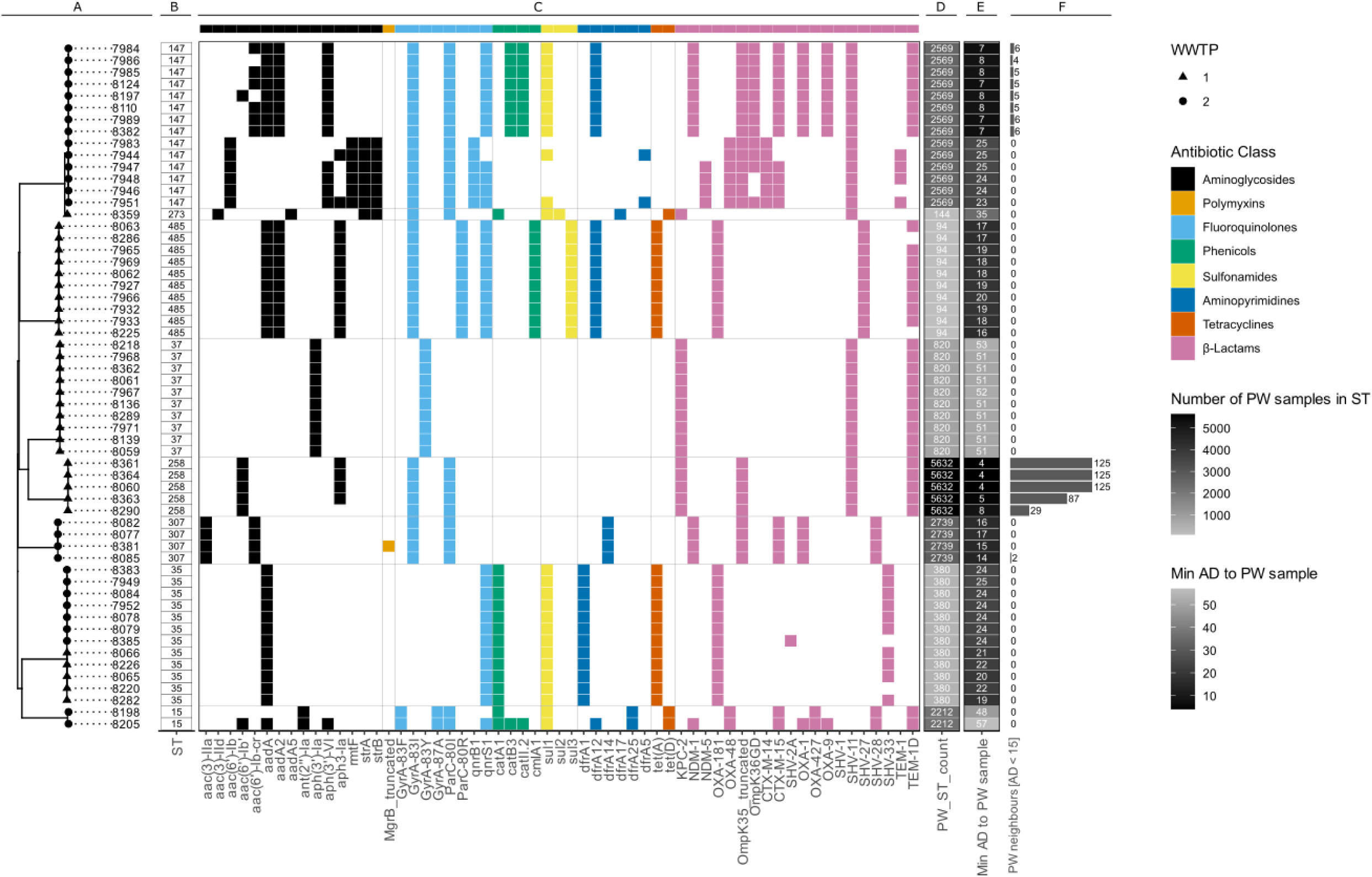
WGS AMR profiling of 58 CP-KP isolates from two WWTPs. A) A phylogenetic tree of 58 wastewater isolates from WWTP 1 and 2. B) Sequence types (STs). C) The presence of ARGs in the isolate assemblies, color-coded by the antibiotic class to which they confer resistance. D) The number of publicly available samples in Pathogenwatch belonging to the same ST as the focal sample. E) The minimal cgMLST allelic distance to the closest publicly available Pathogenwatch genomes. F) The number of publicly available genomes from Pathogenwatch with a core genome MLST (cgMLST) allelic distance (AD) of fewer than 15 alleles from the corresponding wastewater isolate.

Closely related Pathogenwatch genomes, with an allelic distance (AD) of less than 15 to the genomes of wastewater isolates, were predominantly associated with STs that are more frequently observed in the Pathogenwatch dataset. Even though only one out of four isolates belonging to ST307 had two Pathogenwatch neighbors with an AD < 15, all WGS of isolates illustrated low minimal ADs (14-17) to their closest publicly available sample in the Pathogenwatch dataset. Genomes of ST147 wastewater isolates associated with *bla*_NDM-1_ showed up to six closely related Pathogenwatch samples with AD < 15 and minimal ADs as low as seven alleles, while the ST147 isolates associated with *bla*_OXA-48_ and *bla*_NDM-5_ showed greater distance to publicly available samples, with a minimal AD of 23. For ST258, the minimal AD to Pathogenwatch genome entries ranged between 4 and 8. Such close matches were also reflected in the number of Pathogenwatch genome entries with AD < 15, which ranged from 29 to 125. The threshold of an AD < 15 did not show any hits with Pathogenwatch genomes for the other STs, for which the minimal AD to database genomes were instead as follows: 48-57 (ST15), 19-25 (ST35), 51-53 (ST37), 35 (ST273), and 16-20 (ST486) (Figure 4).

It is noteworthy that most Pathogenwatch genomes closely related to the wastewater isolates from this study were from Germany (Figure S2). Moreover, most hits are from fecal isolates, followed by clinical and human-associated samples (Figure S3).

### 3.5 Tracing temporal ST occurrence and SNP dynamics across the sample set

A detailed comparative whole-genome SNP analysis was performed in order to investigate genomic distances between isolates assigned to a particular ST and to facilitate further tracing of putative temporal dynamics across 13 weeks of sampling (Figure 5). A further analysis, including isolates sharing the ST35 background present in samples of WWTP 1 and 2, is available in Figures S4 and S5.

**Figure 5.**
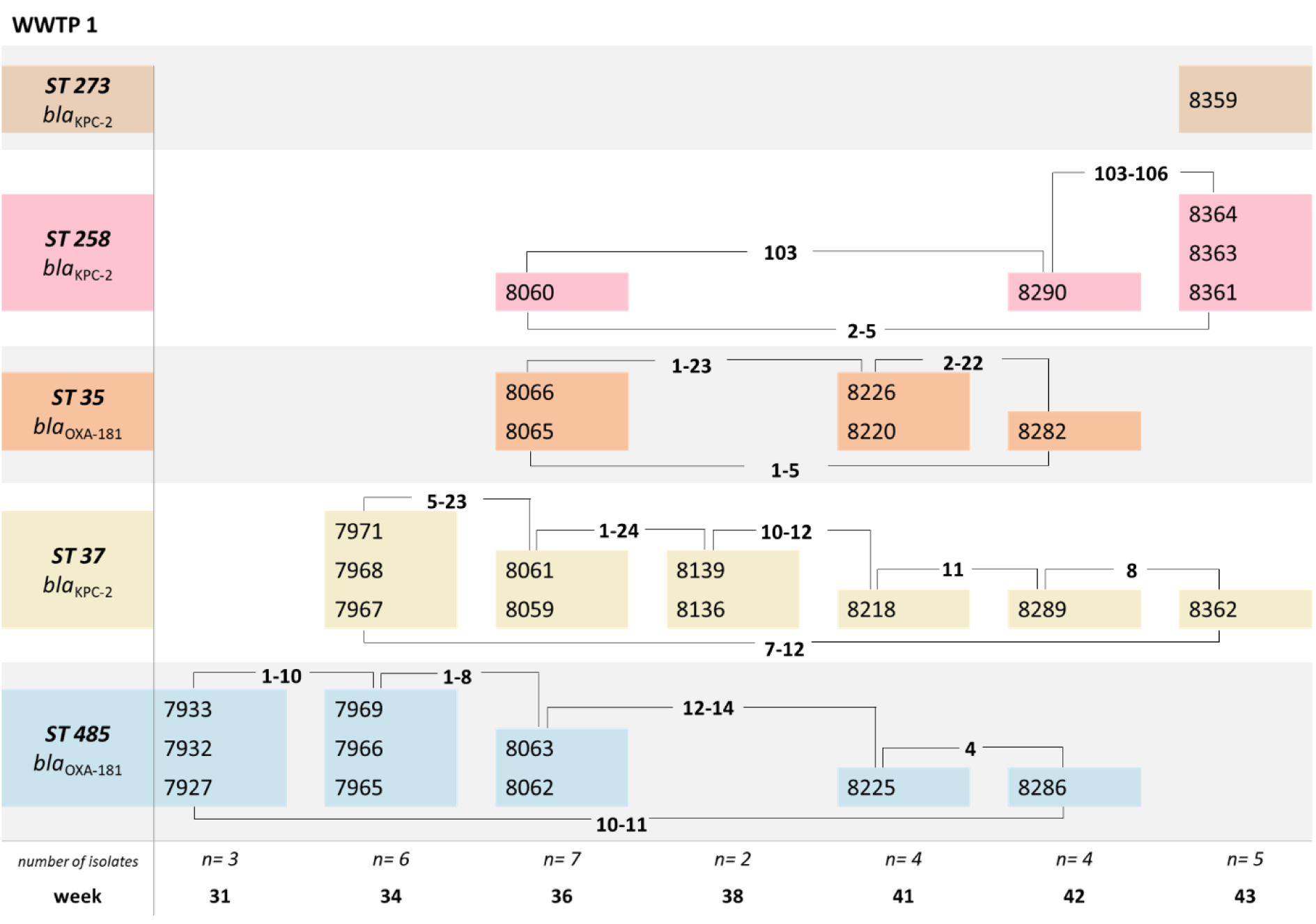
Occurrence, temporal variation, and range of SNP distances of CP-KP WGS per ST detected in WWTP 1. Color-coded STs are assigned to the wastewater sample (week of isolation) indicated on the x-axis, along with the number of isolates subjected to WGS per week. Four-digit strain IDs represent the individual isolates (Figures 4 and S4, Table S4). Single-nucleotide polymorphism (SNP) distances between subsets of isolates and/or sampling weeks are indicated with bold numbers (see Figure S5 for full overview of pairwise SNP distances between isolates). Abbreviations: WWTP, wastewater treatment plant; ST, sequence type; WGS, whole genome sequencing; SNP, single nucleotide polymorphism

Closely related ST485 isolates were detected in WWTP 1, with a genomic distance range between one and 14 SNPs from week 31 to 42. Samples belonging to ST37 showed a minimum genomic distance of 7 SNPs between isolates from weeks 31 and 43, indicating limited genomic diversity within this sequence type across the sampling period. The range of differences between isolates obtained on a specific sampling day is two (week 38) to 21 (week 36). Genomes assigned to ST35 were detected at multiple time points in both WWTPs (Figures 6 and 7). SNP distances revealed distinct clusters among WWTPs differing from each other by at least 14 SNPs (see Figure S5). A group of closely related ST35 isolates obtained from WWTP 1 samples showed a genomic distance range of 1-24 SNPs. One sample (8220, WWTP 1, week 41) exhibited a minimum difference of 19 SNPs to other genomes from WWTP 1, and a minimum difference of 32 SNPs to genomes from WWTP 2 (see Figure S5). The range of SNP differences between isolates assigned to ST258 is 2 to 106 SNPs, with isolate 8290 (week 42) being an outlier (see Figure S5).

**Figure 6.**
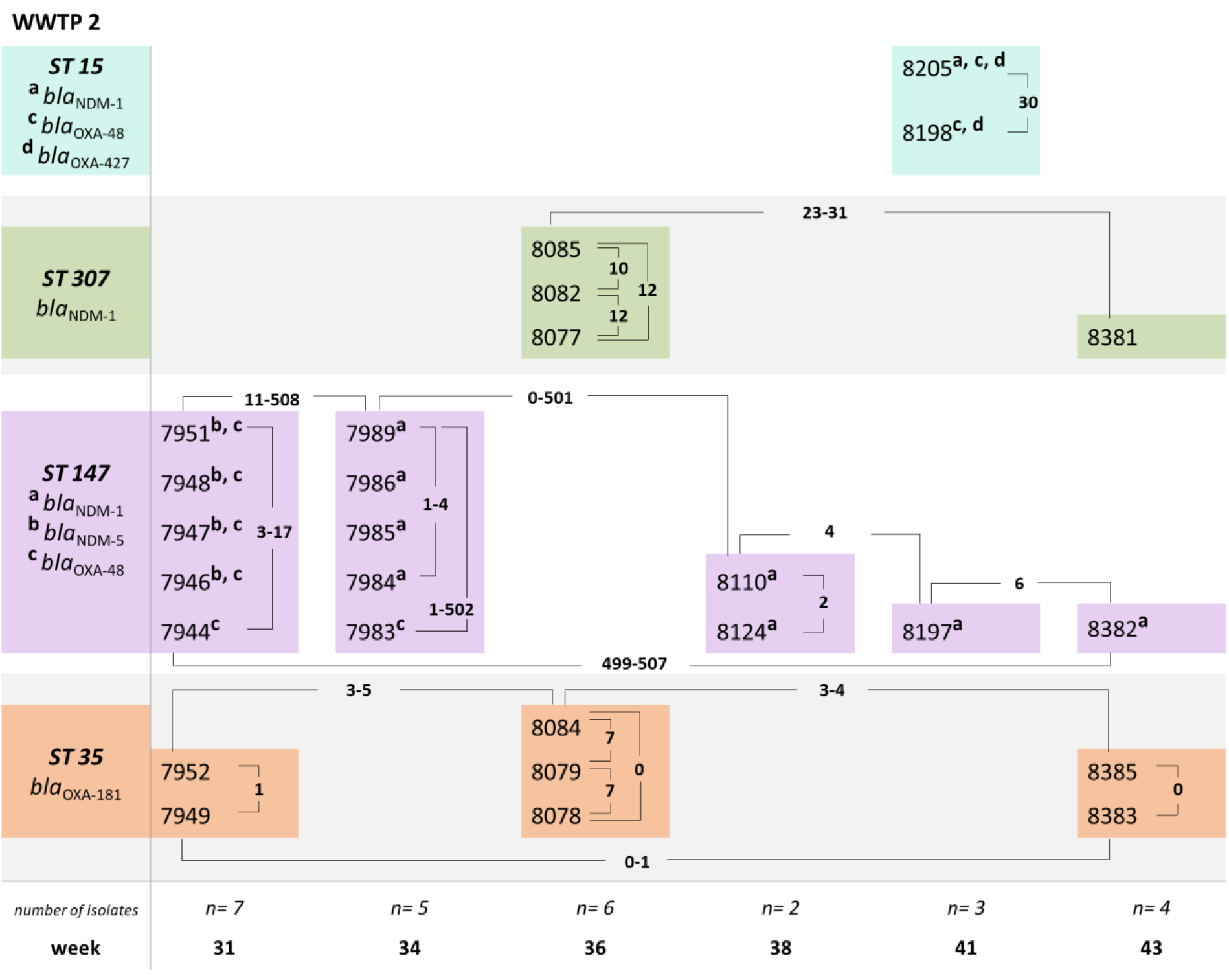
Occurrence, temporal variation, and range of SNP distances of CP-KP WGS per ST detected in WWTP 2. Color-coded STs are assigned to the wastewater sample (week of isolation) indicated on the x-axis, along with the number of isolates subjected to WGS per sampling week. Four-digit strain IDs represent the individual isolates (Figures 4 and S4, Table S4). Single-nucleotide polymorphism (SNP) distances between subsets of isolates and/or sampling weeks are indicated with bold numbers (see Figure S5 for a full overview of pairwise SNP distances between isolates). Abbreviations: WWTP, wastewater treatment plant; ST, sequence type; WGS, whole genome sequencing; SNP, single nucleotide polymorphism

Further investigation of WWTP 2 isolates (Figure 6) revealed that the two ST15 isolates (8198 and 8205 from week 41) differed by 30 SNPs. Genomes assigned to ST307 sampled in week 36 and 43, have 10 to 12 SNPs differences between three isolates from week 36, and a minimum distance of 23 SNPs to the sample 8381 from week 43. Genomes assigned to ST147 showed temporal differences regarding the detection of genomes assigned to two distinct clusters (Figure S5), including different carbapenemase-encoding gene profiles (Figure 5). ST35 genomes obtained from isolates of WWTP 2 seem closely related, with a SNP differences between 0 to 7, regardless of the sampling week.

## 4. Discussion

Here we present a straightforward filtration-based “wet-lab” approach accompanied by WGS analysis to i) quantify and ii) characterize a selected AMR target (CP-KP) in wastewater samples as a proof-of-concept study. Two indicators for wastewater (EC) and AMR (ESBL-EC) viability were used as reference parameters for quantification (iii).

### Quantification of AMR screening target

The workflow was designed based on a lab regime described previously ([54] as cited in [16]) and has been specifically optimized for the detection and quantification of CP-KP. The resulting total verification rate for KP was 93% (297 of 320 identified KEC) including 253 verified CP-KP, which equates to a positive predictive value of 85.1% for CP-KP across all WWTP samples, when using the outlined procedure.

Care should be taken before comparing different approaches aiming at AMR assessment in wastewater samples due to divergent study designs, target choices and abundances. Here, the workflow was specifically amended to target CP-KP. However, CP-E and other carbapenemase-producing gramnegative bacteria isolated from wastewater samples have been quantified before. Thus, findings from previous studies may be useful for broadly classifying procedure performance. In this study, each of the 33 wastewater samples from four distinct WWTPs tested positive for CP-KP. Previous studies reported CP-KP detection at least once in eight out of ten WWTPs in Finland using the same selective agar (25/89 samples tested positive in total = 28.1%) [26]. In comparison, CP-E were isolated from 89 of 100 samples representing 100 urban WWTPs in the Netherlands [16].

The mean colony forming units for CP-KP reported in this study (3.07 x 10³ cfu/100 ml for WWTP 1, 3.79 x 10³ cfu/100 ml for WWTP 2, 7.66 x 10³ cfu/100 ml for WWTP 3 and 1.22 x 10³ cfu/100 ml for WWTP 4) appear slightly higher than values reported in other studies, which often assessed a broader group of species (i.e., CP-E). A previous study across 100 Dutch WWTPs, for instance, reported CP-E values ranging between 0 and 2 × 10^3^ cfu/100 ml, with a mean value of 7.9 × 10^1^ cfu/100 ml [16]. Another study conducted in northern Spain investigating influent samples from two WWTPs, reported mean values ranging between 1 × 10^3^ and 2 × 10^5^ cfu/100 ml CP-E [67]. Presumptive CP-E values obtained for raw wastewater samples of a Japanese WWTP were between 10^3^-10^4^ cfu/100 ml [68]. For Germany, reported figures range between 10^2^ and 10^5^ cfu/100 ml CP-E for influent wastewater of treatment plants not influenced or influenced by hospital wastewater (n= 3/n= 3), respectively. The authors state that the performance of CHROMagar^TM^ ESBL for the quantification and isolation of CP-E is inferior to that of CHROMagar^TM^ mSuperCARBA [69], which was also employed in this study. The low standard deviation values across all WWTPs indicate the reliability and robustness of the target screening procedure established for presumptive CP-KEC and confirmed CP-KP (Figure 2). Moreover, a recent German study suggested that CHROMagar^TM^ mSuperCARBA improves the detection of CP-KEC harboring OXA-48 variants (i.e., OXA-244 and OXA-181) in wastewater samples [69].

Based on the results of our pilot study, the establishment of a structured approach for subsequent method validation according to DIN EN ISO 13843 [70] is required as a next step in advancing the method. This validation includes the categorical performance characteristics (sensitivity, specificity, efficiency, selectivity, false positive rate and false negative rate), determination of the upper limit and consideration of the lower limit of detection, assessment of precision (repeatability and reproducibility), robustness, relative recovery and uncertainty of counting.

### Characterization of a selected AMR target: CP-KP

Following quantification and confirmation of presumptive CP-KP, verified isolates were further phenotypically and genotypically characterized, revealing that a variety of different AST profiles and carbapenemase-encoding genes are associated with the CP-KP wastewater collection.

Since clinical breakpoints suitable for classification of isolates of human or animal origin (as either resistant, intermediate or susceptible) are not applicable to isolates from environmental samples [71], raw MIC values for selected antibiotics representing cephalosporines (e.g., cefotaxime), carbapenems (e.g., imipenem, meropenem), quinolones (ciprofloxacin), aminoglycosides (gentamicin), and sulfonamides (trimethoprim-sulfamethoxazole) are presented in order to facilitate comparability with other studies, especially with respect to One Health integrated surveillance efforts [72].

Interestingly, 14 CP-KP yielded MICs for different carbapenems (Figure 3) below the recommended thresholds for further testing for carbapenemase production [73]. Further characterization of these CP-KP revealed the presence of *bla*_OXA-48_ and/or *bla*_OXA-48_-like genes (e.g., *bla*_OXA-181_). This finding agrees with previous observations for clinical isolates (reviewed in [74, 75]), and wastewater isolates [76]. However, the screening procedure described here detected these phenotypes, since all confirmed CP were subjected to mCIM testing. By applying the screening procedures described in the present study (which involved mCIM testing for all confirmed CP), we were able to detect these phenotypes which might have been missed otherwise. Thus, our approach allows for the detection of a broad range of different genotype/phenotype combinations among CP-KP populations, which is also reflected by the within-sample heterogeneity of the wastewater CP-KP (Figure 4, Figure 5).

Besides carbapenems, most CP-KP isolates yielded MICs above clinical breakpoints for cefuroxime (second generation cephalosporines), cefotaxime and ceftazidime (third generation cephalosporines), sulfonamide and antifolate classes (sulfamethoxazole and trimethoprim combination) and fluoroquinolones (ciprofloxacin), indicating the presence of a broad range of resistance-encoding genes.

### CP-KP representing two different WWTPs include STs of global concern

Unless the entry pathway is exclusive, it is generally not appropriate to draw conclusions about a single source from communal wastewater samples. Unlike samples subjected to microbiological diagnostics from well-defined origins, such as human, food, animal, or process-hygiene sources, this complex matrix comprises multiple concurrent entries. The phenotypical and genotypical diversity of a specific bacterial species obtained from wastewater screening procedures is difficult to predict, as it can vary between sewage treatment plants, sewer systems, the population connected to the system, industrial discharges and sampling days.

A total of 8 different STs, i.e., ST147 and ST273 (both: clonal group 147), ST258, ST35, ST15, ST37, ST307, ST485 were identified in this study. Due to the limited number of isolates subjected to WGS and the lack of long-term data, discussion on distinct ST frequency occurrences is out of scope. Here, WGS was primarily used as a complementary method to confirm the reliability of the wet lab protocol targeting CP-KP and to capture overall CP-KP diversity within individual samples across the sampling period. More research on the subject, especially comparison with metagenomic data, will provide insights about the ability of the proposed approach to capture phylogenetic diversity of CP-KP. Multiple STs (up to four) were identified across the majority of the wastewater samples. Indicating that the presented approach allows for simultaneous detection of genetically diverse CP-KP (Figures 4, 5 and 6). At the time of analysis, there was significant variation in sample availability among different STs. For instance, fewer than 100 samples were available for some STs, such as ST485, whereas others, like ST258, had more than 5,600 samples.

Besides being identified in wastewater samples of this study, CP-KP belonging to ST15 [77], ST147 (and ST273) [77, 78], ST35 [79], ST307 [80], ST258 [81] seem of general clinical importance, whereas isolates belonging to ST485 are less frequently reported [82-84]. Except for ST35, all detected STs have been previously described in studies dealing with influent wastewater samples: ST15 [26, 77], ST37 [26], ST147 [85, 86], ST258 [86, 87], ST273 [88], ST307 [26, 89], ST485 [87]. Even though KP belonging to ST35 were previously detected in wastewater, the respective isolates were negative for carbapenemase production [87, 90].

Some of the CP-KP STs (e.g. ST258, ST307) have been recognized as high-risk clones in human hospitals that are associated with a broad range of ARGs [80, 91, 92]. CP-KP ST258, for instance, is of public health concern because of its widespread distribution, its frequent multidrug resistance, and its ability to rapidly exchange plasmid borne resistance determinants with other *Enterobacteriaceae* ([93] and reviewed in [94]). It has been discussed that adaptive mutations likely contribute to ST258 persistence among hospitalized patients [95], and mutations have been detected among a single patient colonized by CP-KP over a 4.5-year period [96]. Thus, long-term enteral colonization of humans, either in terms of hospitalized patients or even among the general population, needs to be considered as a possible source for CP-KP in wastewater samples. Furthermore, environmental sources need to be addressed as well. Closely related CP-KP genomes belonging to ST147 have been isolated from different sampling sites influenced by hospital wastewater. These sites include a path from patient rooms (toilets and showers) to the WWTP effluent. In particular, the limited genomic diversity together with the stable presence of the *bla*_NDM-1_ (often in association with *bla*_OXA-48_) gene among all isolates investigated was discussed as a sign for long-term persistence of this particular clone in the respective sewage system [97]. The two clusters of ST147 genomes from isolates of WWTP 2 in the present study carry either *bla*_OXA-48_ with or without *bla*_NDM-5_ genes, or *bla*_NDM-1_. Genome distances of >500 SNPs (Figures 6 and S5), indicate the presence of at least two different clusters, which were detected simultaneously in week 34.

Of note, one isolates (8205) belonging to ST15 revealed the presence of three distinct carbapenemase-encoding genes (*bla*_OXA-48_, *bla*_OXA-427_, *bla*_NDM-1_). Such accumulation of resistance genes appears to be rare, but not unprecedented, as reported for a clinical ST147 isolate [98].

Comparative analysis of samples across the wastewater network based on 16S metagenomic sequencing revealed that bacterial ß-diversity was more similar between WWTP influent samples and samples from a community wastewater collection point than between WWTP influent samples and hospital wastewater samples. This finding suggests the potential to detect relevant, and potentially novel pathogen-resistance phenotypes within the population connected to the system [99].

### Phylogenetic relationship between WGS

So far, only limited genomic diversity has been reported for CP-KP isolates from wastewater samples [69], hospitalized patients [95], and hospital environments [79], even from sampling campaigns spanning several years. These findings raise questions regarding the expected mutation rates of CP-KP in its natural habitat as part of the gut-colonizing microbiota. Additional uncertainty exists regarding mutation rates in more challenging environments, such as in patients receiving treatment (antibiotics) or in the wastewater system [97].

Of note, the majority of the Pathogenwatch samples closely related to wastewater isolates from this study are submissions from Germany. These samples are mostly of fecal origin, followed by clinical and other human-associated sources (Figure S2-S3), indicating the influence of the population connected to the wastewater system on the composition of bacterial load in influent wastewater [100, 101].

When analyzing the phylogenetic relationship or clonality of KP using SNPs, the expected mutation rate for that particular species is an important parameter. Current literature provides, due to different methods i.e., core genome (cg), maximum common genome (mcg), and study goals, a variety of mutation rates for KP, including CP-KP and different STs [102]. Recently, genome-wide nucleotide substitutions of multidrug-resistant KP sharing the clonal group 147 (i.e., ST147, ST273 and ST392) revealed an evolutionary rate range from ∼1x10⁻⁶ to 2x10⁻⁶ substitutions/site/year (≈ 3–8 SNPs/genome/year) [78]. The genomes presented here span a sampling period of 13 weeks, thus a range of 0.75 to 2 SNPs per genome seems a reliable estimation for clonally related isolates. A further study on pairwise SNP distances between CP-KP ST35 outbreak strains in Germany showed very limited diversity within clonally related clusters (i.e., 0 to 6 SNPs) over a period of 30 weeks [79], suggesting careful interpretation of SNP distances between genomes from wastewater-associated isolates.

Taken together, further research and development efforts are required to increase our ability to trace back the source of CP-KP in wastewater samples and to address significant knowledge gaps regarding survival and residence of clinical important CP-KP in wastewater systems, including biofilms. However, based on the Pathogenwatch matches, feces from enteral colonized people appear to be a plausible source of the isolates reported in this study. Nevertheless, for reliable conclusions about the ultimate sources of CP-KP from wastewater additional research is needed. The inclusion of the wastewater pathway may provide early insight into the emergence of novel pathogen-resistance combinations and thus could further strengthen integrated genomic surveillance efforts and public health response.

### Reliability indicators for wastewater (EC) and AMR (ESBL-EC)

Regarding overall ranges of cfu/100 ml concentration levels, EC and ESBL-EC counts showed no explicit temporal variation while the observed values were generally consistent with those reported in studies from various European countries [16, 17, 29, 103-105], North America [106], and Africa [107]. This consistency indicates the suitability of these indicators as proxy for fecal load and AMR in wastewater samples, respectively. The WHO Tricycle Protocol has previously stated that ESBL-EC is a relevant and representative proxy for AMR [27]. Including independent quality control indicators is therefore essential to ensure sample validity and for to enable comparable quantification of target organisms.

A recent study compared culture-based detection with real-time (RT)PCR quantification of pathogens in wastewater samples and demonstrated that RT-PCR based depends strongly on primer specificity and the presence of conserved target regions. In that study, two of six target species were not detected by RT-PCR [108]. The authors attributed this limitation to genetic variability in environmental strains, including mutations in primer-binding regions or the absence of specific housekeeping or virulence genes, which can result in non-amplification despite the presence of viable cells. The study further demonstrated that integrating culture-based and molecular methods improved detection sensitivity and provided complementary insights [108].

### Study limitations

This study was designed and implemented as a proof-of-principle approach, confirming its ability to detect different CP-KP in wastewater, and the procedure was rigorously tested within the scope of this study, but has several important limitations. Those include the restricted geographic coverage and number of WWTPs included, the putative process bias associated with the use of filter membranes used for bacterial concentration, and the possible loss of clinically relevant CP-KP isolates due to the chosen incubation regimen and selective media. A validation process according to international standards is required. These limitations need to be addressed in subsequent studies to further improve targeted CP-KP detection in wastewater samples.

### Outlook

In the absence of ‘gold standards’, the assessment of method performance indicators, including sensitivity and specificity, for AMR quantification in wastewater samples requires a comparative approach that bridges disciplines to ensure reliability of results. While culture-based WGS allows for detailed characterization of individual isolates, including precise identification of ARGs and a deep understanding of phylogeny, it might introduce selection biases [109]. Such selection bias can be addressed by incorporating culture-independent metagenomic approaches alongside culture-based methods. While sequencing depth and related parameters are critical for detecting rare genetic determinants, metagenomic data from the same samples enable direct comparison between culture-based and culture-independent results and provide complementary insights into the diversity and abundance of ARGs and ARBs in the wastewater system [110]. Future studies should continue to investigate the complementary nature of isolate-based sequencing and metagenomic analyses to assess the broader resistome in wastewater. However, as previously shown in specific settings, it is much more feasible to monitor a single ARB target than an assortment of AMR pathogens and genes [51] and the importance of sensitive methods for targeting rare subpopulations such as CP-KP was addressed only recently [30].

Taken together, this proof-of-principle approach demonstrates the feasibility of detecting CP-KP in wastewater. In the context of regulatory requirements, the availability of validated and robust methods is essential to ensure comparability and reliability within a regulatory framework. Therefore, a validation process that adheres to international standards is imperative.

## Supporting information

Werner-et-al-2026_Supplementary-Material

## Acknowledgements

We sincerely appreciate the support of the wastewater treatment plant operators and their employees participating in AMELAG, with special thanks to the wastewater disposal companies in Mecklenburg-Vorpommern - Nordwasser GmbH (Kläranlage Rostock), Schweriner Abwasserentsorgung Eigenbetrieb der Landeshauptstadt Schwerin, Neubrandenburger Wasserbetriebe GmbH, and Abwasserwerk Greifswald, Eigenbetrieb der Universitäts-und Hansestadt Greifswald. In addition, we would like to thank the Sequencing Core Facility of the Genome Competence Center of the Robert Koch Institute (RKI) for providing excellent sequencing services.

We thank all Members of the AMELAG team and highly acknowledge Ulrike Braun and Markus Lucas for project coordination, and the organization of sample logistics. Special thanks go to the involved authorities.

## Data availability

The Illumina sequencing data generated in this study have been deposited in the European Nucleotide Archive (ENA) under the BioProject accession number PRJEB107251.

## Contributions

KW, AB, MH and BW designed the project. KW, MH, CB, BV, and BW conceived and designed the experiments. KW, DB, JB, KL, and AB performed laboratory analysis and isolate culturing. CB and SS prepared libraries and sequenced isolates. KW, BW, VB, AB, SAW, CB, CF and MH analyzed the data. KW and BW wrote the first draft. All authors have read and approved the final draft of the manuscript.

