## Supplementary material for "Bridging disciplines towards wastewater-based surveillance of antimicrobial resistance: frequency, local dynamics, and genomic characteristics of carbapenemase-producing *Klebsiella pneumoniae*": Werner-et-al-2026_Supplementary-Material

**S1. Pretest results: evaluation of two different temperature regimes for detection of KP on chromogenic media**

Two temperature regimes were initially evaluated with respect to their effect on detection frequencies of the primary target species (CP-KP) as well as fecal and AMR indicators (EC, ESBL-EC) on chromogenic media. In total, six influent wastewater samples obtained over a period of three consecutive weeks were analyzed, with up to 30 subcultures per target and incubation regime identified to species level. Median values obtained for confirmed KP were 15% (37 °C for 18h) and 65% (36 ± 1 °C for 4h followed by 21h at 44 ± 0.5 °C as described by [1] as cited in [2]) (Figure 2). Differences for confirmed KP were highly significant ($p < 0.001$). EC and ESBL-EC exhibited a non-significant difference following species verification with median values for i) EC of 95% (37 °C) / 100% (44 °C) and ii) ESBL-EC of 93% (37 °C) / 100% (44 °C). More details are provided in supplemental Table S3.

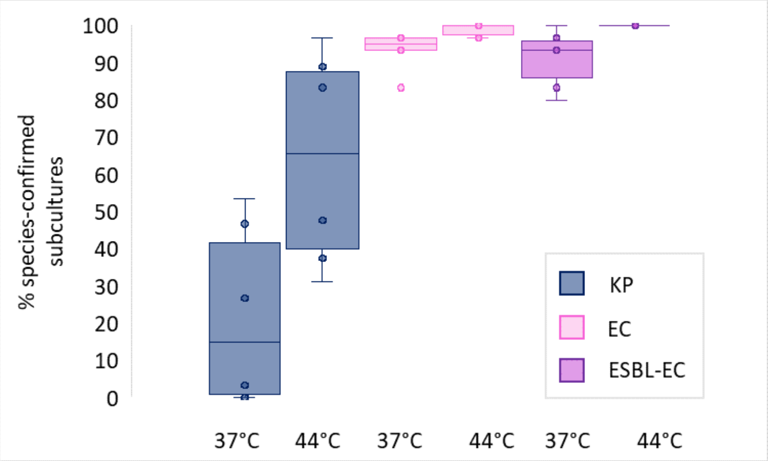


***Figure S1. Colony count results of two different temperature regimes.***

*Comparative assessment of two different temperature regimens regarding target species by orders of magnitude obtained from colony counts of wastewater-loaded filters on selective media (CHROMagar™ mSuperCARBA (KP), CHROMagar™ Orientation (EC), and CHROMagar™ ESBL (ESBL-EC)). Up to 30 subcultures per target and incubation regime were analyzed from six individual wastewater samples. Boxplots indicate the percentage [%] of species-confirmed subcultures of total isolates analyzed.*

*Abbreviations: KP,* Klebsiella pneumoniae*; EC,* Escherichia coli*; ESBL-EC, extended-spectrum beta-lactamase* E. coli.

Pretesting of different incubation protocols revealed significantly higher detection rates for the target organisms after incubation at 44 °C compared to 37 °C, incubation at elevated temperatures inhibits off-target-growth, including gram-negative non-fermenters such as *Aeromonas* spp. [3, 4]), as well as other species of the KEC group, as shown previously for *K. oxytoca* and *Enterobacter cloacae* [5]. This allows for more precise target colony enumeration due to less unspecific growth on the respective filters. Thus, the 44 °C-incubation regime is suitable to detect a low abundance target (CP), as described previously regimes ([1] as cited in [2]). However, more studies are necessary to assess putative loss of temperature-sensitive strains due to elevated incubation temperatures [6, 7].

***Table S1. PCR primers for carbapenemase-encoding genes***

| Target gene | Sequence [5'-3'] | $T_M$ [°C] | Amplicon size [bp] | Reference |
|---|---|---|---|---|
| *bla*$_{IMP}$ (all genes) | F: CATGGTTTGGTGGTTCTTGT | 55.3 | 447 | Gröbner et al. (2009) [8] |
| | R: ATAATTTGGCGGACTTTGGC | 55.3 | | |
| *bla*$_{KPC}$ (all genes) | F: CAGCTCATTCAAGGGCTTTC | 57.3 | 533 | Gröbner et al. (2009) [8] |
| | R: AGTCATTTGCCGTGCCATAC | 57.3 | | |
| *bla*$_{NDM}$ | F: CTGAGCACCGCATTAGCC | 58.2 | 754 | Pfeifer et al. (2011) [9] |
| | R: GGGCCGTATGAGTGATTGC | 58.2 | | |
| *bla*$_{OXA-48}$-related genes | F: AAATCACAGGGCGTAGTTGTG | 57.9 | 555 | Gröbner et al. (2009) [8] |
| | R: GACCCACCAGCCAATCTTAG | 59.4 | | |
| *bla*$_{VIM}$ | F: GTTTGGTCGCATATCGCAAC | 57.3 | 384 | Pitout et al. (2005) (modified) [10] |
| | R: CGAATGCGCAGCACCRGGAT | 62.4 | | |

*Primer pairs used for PCR of carbapenemase-encoding genes bla$_{IMP}$, bla$_{KPC}$, bla$_{NDM-1}$, bla$_{OXA-48}$ and bla$_{VIM}$. PCR was performed in 25 µL reactions containing: 1x reaction buffer (New England Biolabs, Germany), 120 µM dNTP mix (New England Biolabs, Germany), 0.2 µM of each primer (Eurofins, Germany), 0.05 µl Taq DNA polymerase (New England Biolabs, Germany), and 2.5 µl template. Thermal cycling was carried out at 94 °C for 3 min, followed by 30 cycles of [94 °C for 30 s, 55 °C for 20 s, 72 °C for 45 s], and a final extension step at 72 °C for 5 minutes. Template preparation for PCR was conducted by heat lysis of single colonies, cultivated overnight on columbia blood agar plates (Thermo Scientific, UK). PCR products were visualized on 1.5% agarose-TAE gels stained with ROTI®GelStain (Roth, Germany) by means of the Azure 300 imager (Azure Biosystems, USA). A 100 bp DNA ladder (New England Biolabs, Germany) was used for size comparison.*

*Abbreviations: F, forward; R, reverse; TM, melting temperature*

***Table S2. Pathogenwatch samples for different* Klebsiella pneumoniae *sequence types***

| Sequence Type (ST) | Number of samples on Pathogenwatch |
|---|---|
| ST258 | 5632 |
| ST307 | 2739 |
| ST147 | 2569 |
| ST15 | 2212 |
| ST37 | 820 |
| ST35 | 380 |
| ST273 | 144 |
| ST485 | 94 |
| **Total** | **14590** |

*Summary of the number of publicly available* Klebsiella pneumoniae *genomes on Pathogenwatch for each sequence type (ST) identified in the wastewater isolates, as of February 20, 2025.*

***Table S3**. **Comparison of two incubation regimes for the detection of Carbepenmase-producing*** **Klebsiella pneumoniae, Escherichia coli*****, and ESBL-*****E. coli*****.**

| **incubation temp. [°C]** | **sample** | **KEC including KP** | | | | | **EC** | | | **ESBL-EC** | | |
|---|---|---|---|---|---|---|---|---|---|---|---|---|
| | | subcultures | KEC (confirmed) | KP (confirmed) | % (KEC) | % (KP) | subcultures | species confirmed | % | subcultures | species confirmed | % |
| **37** | 1 | 30 | 12 | 8 | **40.0** | **26.7** | 30 | 28 | **93.3** | 30 | 28 | **93.3** |
| | 2 | 30 | 5 | 1 | **16.7** | **3.3** | 30 | 29 | **96.7** | 30 | 24 | **80.0** |
| | 3 | 15 | 3 | 0 | **20.0** | **0.0** | 30 | 25 | **83.3** | 30 | 29 | **96.7** |
| | 4 | 30 | 17 | 16 | **56.7** | **53.3** | 30 | 29 | **96.7** | 30 | 25 | **83.3** |
| | 5 | 30 | 20 | 14 | **66.7** | **46.7** | 30 | 29 | **96.7** | 30 | 30 | **100** |
| | 6 | 30 | 4 | 0 | **13.3** | **0.0** | 30 | 28 | **93.3** | 30 | 28 | **93.3** |
| **44** | 1 | 27 | 27 | 24 | **100** | **88.9** | 30 | 30 | **100** | 30 | 30 | **100** |
| | 2 | 16 | 16 | 5 | **100** | **31.3** | 30 | 30 | **100** | 30 | 30 | **100** |
| | 3 | 21 | 21 | 10 | **100** | **47.6** | 30 | 30 | **100** | 30 | 30 | **100** |
| | 4 | 30 | 30 | 25 | **100** | **83.3** | 30 | 29 | **96.7** | 30 | 30 | **100** |
| | 5 | 30 | 29 | 29 | **96.7** | **96.7** | 30 | 30 | **100** | 30 | 30 | **100** |
| | 6 | 8 | 8 | 3 | **100** | **37.5** | 30 | 29 | **96.7** | 30 | 30 | **100** |

*Results of the comparison of two incubation protocols regarding the isolation of target species in wastewater samples. Target species were suspect carbapenemase-producing* K. pneumoniae *(CP-KP), total* E. coli *(EC), and ESBL-*E. coli *(ESBL-EC). The trial comprised influent wastewater samples from two WWTPs, respectively, in three consecutive weeks, i.e., six samples in total. For each target, two volumes of wastewater were filtered in triplicates, resulting in six agar plates per target and incubation regime. If possible, five colonies per plate were used for species-confirmation by MALDI-TOF MS, resulting in a maximum of 30 colonies analyzed per incubation regime. Percentages [%] indicate the ratio of species-confirmed isolates of total isolates analyzed.*

*Abbreviations: temp., temperature; KEC,* Klebsiella, Enterobacter, Citrobacter*; KP,* Klebsiella pneumoniae*; EC,* Escherichia coli*; ESBL, extended-spectrum beta-lactamase*

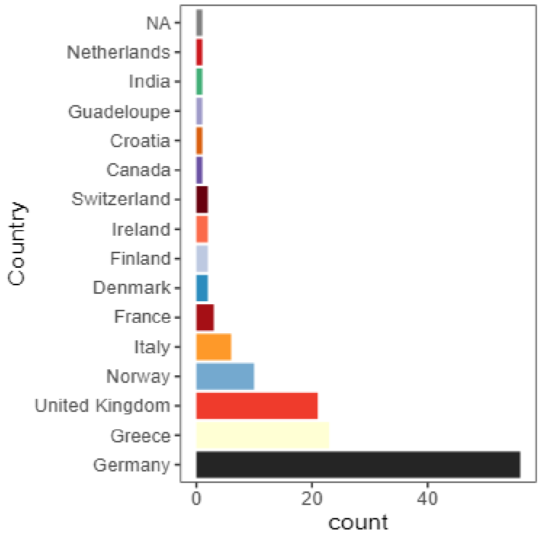


***Figure S2. Contributing countries to* Klebsiella pneumoniae *samples on Pathogenwatch.*** *Number of publicly available Pathogenwatch samples with a cgMLST allelic distance of fewer than 15 alleles to at least one of the 58 wastewater isolates from this study, grouped by country of origin (n= 133). Colours indicate countries.*

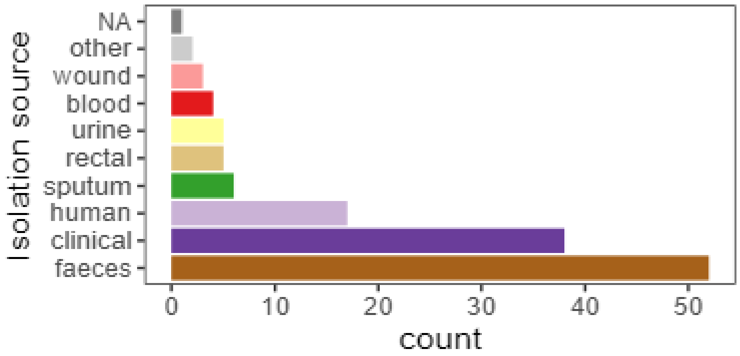


***Figure S3. Isolation sources of* Klebsiella pneumoniae *samples on Pathogenwatch.*** *Number of publicly available Pathogenwatch samples with a cgMLST allelic distance of fewer than 15 alleles to at least one of the 58 wastewater isolates from this study, grouped by isolation source (n= 133). Colours indicate isolation sources.*

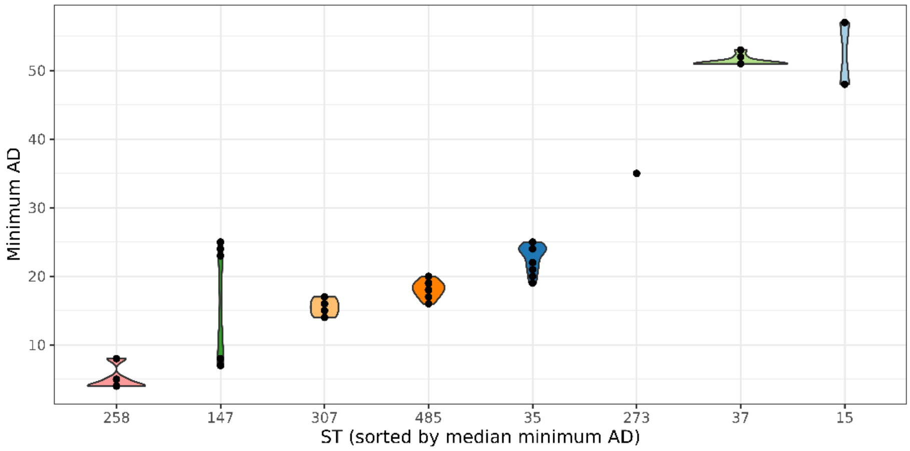


***Figure S4. Distribution of minimal cgMLST allelic distances between wastewater isolates and Pathogenwatch samples.*** *Violin plots showing the distribution of minimal cgMLST allelic distances (AD) between wastewater isolates from this study and publicly available Pathogenwatch samples, grouped by sequence type (ST). Each dot represents the minimal AD for a given wastewater sample.*

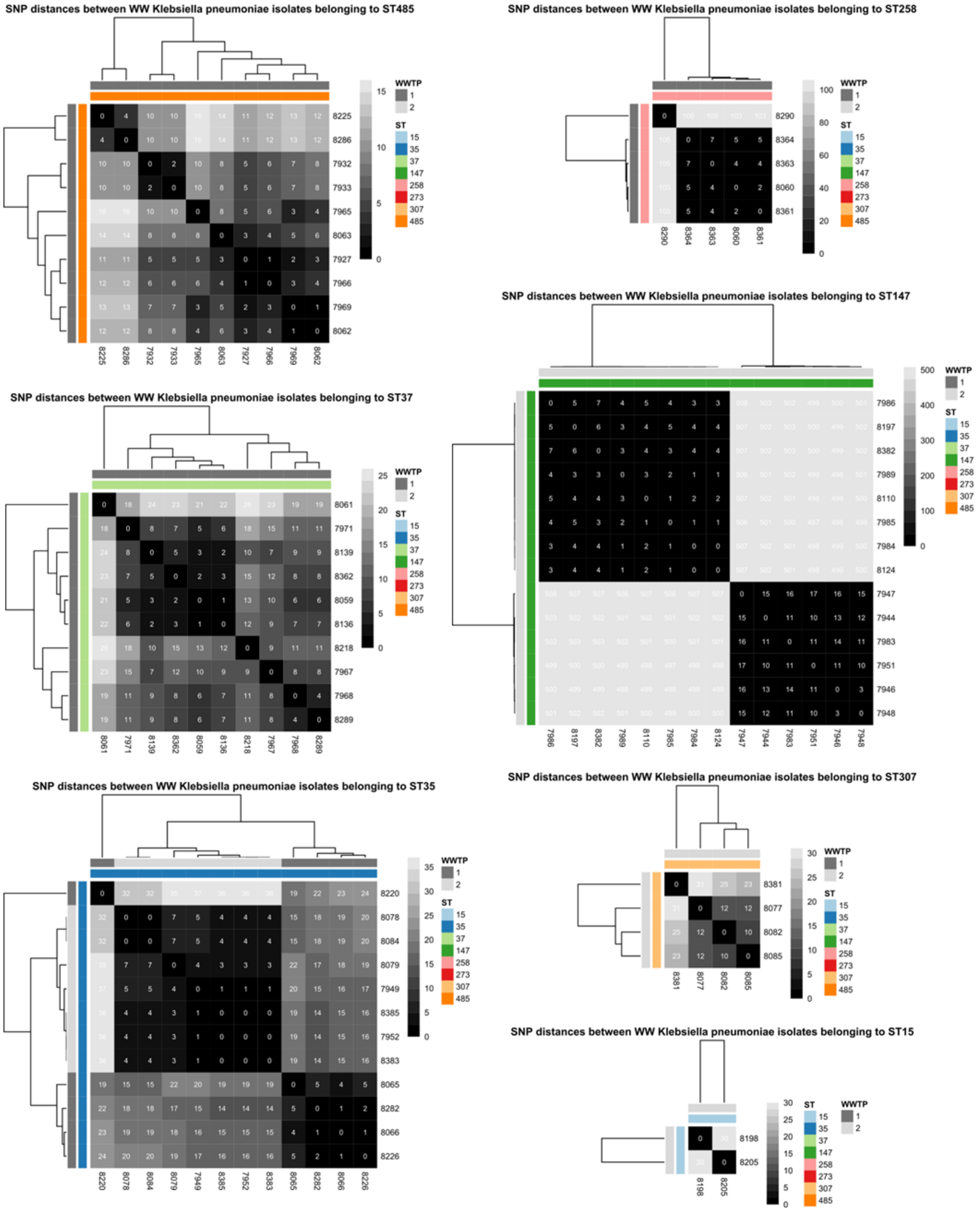


***Figure S5**. **SNP distances of WGS data of*** **Klebsiella pneumoniae** ***isolated from two WWTPs.*** *Heatmap of pairwise single nucleotide polymorphism (SNP) distances among* Klebsiella pneumoniae *wastewater isolates, based on snippy analysis. Reference genomes were samples 8286 (ST485), 8289 (ST37), 8383 (ST35), 8290 (ST258), 7985 (ST147), 8082 (ST307), and 8198 (ST15).*
